# Melanopic stimulation does not affect psychophysical threshold sensitivity for luminance flicker

**DOI:** 10.1101/2021.07.08.451597

**Authors:** Joris Vincent, Edda B. Haggerty, David H. Brainard, Geoffrey K. Aguirre

## Abstract

In addition to the cone photoreceptors the retina contains intrinsically photosensitive retinal ganglion cells (ipRGCs). These cells express the photopigment melanopsin and are known to be involved in reflexive visual functions such as pupil response and photo-entrainment of the circadian rhythm. It is possible that the ipRGCs contribute to conscious visual perception, either by providing an independent signal to the geniculo-striate pathway, or by interacting with and thus modifying signals arising from “classical” retinal ganglion cells that combine and contrast cone input. Here, we tested for the existence of an interaction by asking if a 350% change in melanopsin stimulation alters psychophysical sensitivity for the detection of luminance flicker. In Experiment 1, we tested for a change in the threshold for detecting luminance flicker in three participants after they adapted to backgrounds with different degrees of tonic melanopsin stimulation. In Experiments 2 and 3, this test was repeated, but now for luminance flicker presented on a transient pedestal of melanopsin stimulation. Across the three experiments, no effect of melanopsin stimulation upon threshold flicker sensitivity was found. Our results suggest that even large changes in melanopsin stimulation do not affect near-threshold, cone-mediated visual perception.

## Introduction

Under daylight conditions, signals originating in the cone photoreceptors pass through several classes of retinal ganglion cells to support visual perception. Also active in daylight are the melanopsin-containing, intrinsically photosensitive retinal ganglion cells (ipRGCs). The ipRGCs mediate numerous physiologic effects of light, including variation in pupil size and photoentrainment of the circadian rhythm (Berson et al., 2002; Lucas et al., 2001). Beyond these “reflexive” visual functions, several studies have examined if signals from the ipRGCs contribute to visual perception. When human observers are presented with stimuli that include an increase in melanopsin stimulation, participants report that the spectral change appears as increase in “brightness” (Brown et al., 2012; Spitschan, 2019; Zele et al., 2018a; Zele et al., 2019; Allen et al., 2017; Allen et al., 2019; Yamakawa et al, 2019; DeLawyer et al., 2020). Studies of this kind support the claim that ipRGC signals have a direct effect upon perception.

In addition to this direct effect, signals from the ipRGCs may interact with those carried by the cones, producing effects upon perception by an indirect mechanism (Figure 1a). Such interactions could occur within the ipRGCs themselves, as these cells receive signals from the cones. In addition, in both the primate and rodent retina, a subset of ipRGCs send recurrent axon collaterals to the inner plexiform layer (Joo et al., 2013) where they are hypothesized to influence the sensitivity of cone inputs to the “classical” RGCs (i.e., midget, parasol, small bistratified). This provides another possible site for an indirect effect of melanopsin on signals originating in the cones. Electroretinogram (ERG) recordings in mice support this idea, as reduced b-wave amplitude responses to cone-directed light flashes are observed when presented on a melanopsin-stimulating background (Allen 2014). A third potential point of interaction is present in the projections of the ipRGCs to the lateral geniculate nucleus (LGN), where they signal overall retinal irradiance in both a tonic and phasic manner (Dacey et al., 2005; Brown et al., 2010; Brown et al., 2012; Schmidt et al., 2014). Therefore, signals from the melanopsin-containing ipRGCs could interact with the classical RGC pathways downstream from the retina.

**Figure 1.**
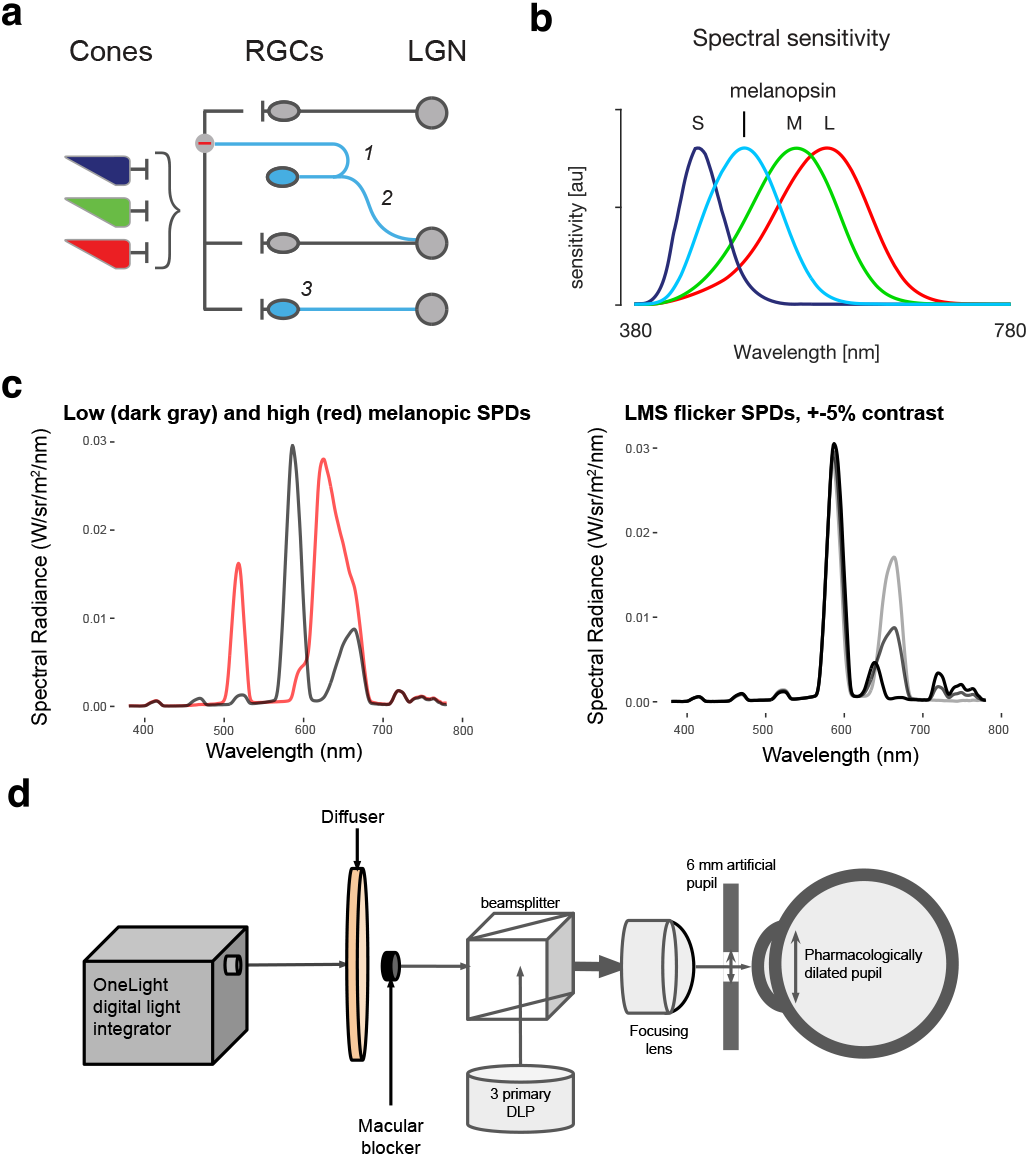
Theoretical background and experimental design. **a)** Retinal wiring schematic. There are several hypothesized locations at which signals from the melanopsin-containing ipRGCs (blue) might interact with signals from the cones, which are predominantly conveyed by the classical RGCs (gray). (1) Recurrent axon collaterals from ipRGCs can provide inhibitory signaling on cone pathways; (2) ipRGCs can project to LGN cells that also receive inputs from the classical RGCs; and (3) cone and melanopsin signals may interact within the ipRGCs themselves. **b)** Spectral sensitivities of the L, M and S cones and of the melanopsin-containing ipRGCs. Each sensitivity is normalized to a peak of unity. **c)** *Left*: Spectral radiance of the low-(dark gray) and high-(red) melanopic backgrounds. *Right*: Example spectral modulations for LMS directed flicker. The dark gray line is the spectral radiance of the low-melanopic background (same as in the panel on the right). The black and light gray lines provide the spectral radiance of the low and high (5% contrast) arms of the 5% LMS directed flicker around this background. **d)** Apparatus schematic. Light produced by the digital light integrator was passed through a randomized optical fiber and lens assembly (not shown) and was imaged on a diffuser, to produce a spatially uniform field. Light from a 3 primary DLP projector was optically admixed through a beam splitter with the light from the digital light synthesizer (lenses and neutral density filters not shown). The participant viewed the combined stimulus through a lens, such that the diffuser subtended 27.5° (diameter), with the central 5° (diameter) macular region occluded.

Here we examine if the threshold for detection of flickering light directed at the luminance channel is altered by modulation of the melanopic excitation produced by the stimulus background. Over three experiments, we measured the threshold at which human observers could detect a 5 Hz modulation of cone contrast. In each experiment, measurements were made under conditions of low-and high-melanopic excitation, with the experiments differing in the timing of the melanopsin modulation, and in the means by which the stimuli were produced. We choose 5 Hz flicker as we expect that this stimulus will predominantly drive the conventional RGCs (e.g., the parasol and midget classes), and less so the relatively sluggish ipRGCs. We find no evidence in our measurements that the change in melanopsin excitation alters the detection of luminance flicker beyond what could be accounted for by imprecision in stimulus generation. These results constrain the ways in which signals from classical and melanopsin-containing RGCs might be theorized to interact.

## Methods

### Overview and participants

Psychophysical sensitivity to flicker directed at the L, M, and S cones was measured across two conditions that differed in melanopic excitation. Measurements were made in three separate experiments. The three experiments were conceptually similar, but differed in the details of the stimuli. A total of six participants contributed data. All participants had normal color vision as assessed by the Ishihara test for color deficiencies. In Experiment 1, the three participants were the three authors of the study (P1, P2, P3), all male, ages 48, 58, and 26 years old, respectively. Three participants naïve to the hypotheses of the study (P4, P5, P6) were also recruited (2 male, ages 29, 25, 23, respectively) and participated in Experiment 2. Experiment 3 obtained data from participants P3, P4, and P5. This study was approved by the University of Pennsylvania Institutional Review Board, and all participants provided written consent. Due to an oversight, participants who were also investigators in the study were not consented in advance of their participation, but each did provide consent *ex post*. This deviation was reported to the Institutional Review Board. The investigators have all consented to the possiblity that their personal data is likely to be identifiable.

### Varying background melanopic excitation

In each experiment, a flicker stimulus was presented around one of two different background light spectra that varied in their melanopic excitation (low or high). The pair of low-and high-melanopic spectra were designed through silent substitution (Spitschan et al., 2015, Estévez & Spekreijse, 1982) to target melanopsin while attempting to silence the L, M and S cones despite their overlapping spectral sensitivities (Figure 1b, c).

Our approach to generating photoreceptor-directed spectral modulations has been described previously (Spitschan et al., 2015, Spitschan et al., 2017). Briefly, cone fundamentals were based on the International Commission on Illumination (CIE) physiological cone fundamentals (CIE 2007). The CIE standard only specifies fundamentals up to field sizes of 10 degrees, so we obtained estimates for our 27.5 degree field by extrapolation of the formula. A 6 mm pupil was assumed. Spectra were tailored to account for the age-predicted lens density for each participant. The melanopic backgrounds and pedestals were generated with a OneLight digital light synthesizer (VISX Spectra), controlled by an Apple Macbook Pro, using custom MATLAB (Mathworks) software. The change from the low-to high-melanopic background constituted a nominal 350% increase melanopsin excitation. In Experiments 1 and 2, the OneLight was also used to produce cone-directed flicker (Figure 1c -right). The luminance of the stimulus field for the background spectrum varied somewhat from session to session but was on the order of 275 cd/m2.

### Stimulus delivery

The spectral output of the OneLight was imaged onto a custom diffuser to produce a circular, uniform field of 27.5 degrees diameter (Figure 1d). Because spatial variation in macular pigment produces variation in photoreceptor spectral sensitivity, the central 5 degrees (diameter) of the field were obscured with black ink. We considered the possibility that scattered light from the stimulus field onto the obscured macular region could confound the measurement. To minimize this, steady light provided by an HP Notebook Companion digital light projector (DLP) was admixed with the optical stimulus from the OneLight (Figure 1e). The admixed light consisted of a central spot overlapping with, and slightly extending past, the central 5 degrees; and an annulus just inside of, and extending outward from, the outer edge of the 27.5 degree stimulus field. The pixels of the DLP aligning with the central blocker and the outer stimulus were set to full-white (RGB input settings = [1 1 1]), and everywhere else to off ([0 0 0]). In the center of this white central region, the DLP output was set to form a small, red cross that participants were instructed to use as a fixation target. The output intensity of the DLP was reduced by passing the projector beam through 3.3-log units of neutral density filter, then imaged onto a custom diffuser screen to form an image, which was then admixed with the optical signal from the OneLight using a beam splitter. The admixed light from the “on” region of the projector image was ∼11 cd/m^2^. In informal observations we found that this was sufficient to render the flicker stimulus undetectable in the central region, even at the highest contrast level used. In sum, the participant viewed the full 27.5 degree annular stimulus region produced by the OneLight stimulus component, with the central 5 degrees and region exterior to the annulus masked by admixed white light.

Spectroradiometric measurements of the stimulus field were taken with the admixed steady light. Measurements integrated over a 1 degree portion of the stimulus, centered at an eccentricity approximately halfway between the obscured macular region and the outer edge of the stimulus field. These measurements were made with the projector light present and absent, and with the OneLight full-power output present or absent. The median spectrum added by the projector light, across conditions, was taken to represent the scatter of the projector light onto the stimulus field. During pilot testing, we found that the projector spectrum contribution to the stimulus region was 2.63 cd/m^2^. This scatter from the admixed steady light from the projector onto the stimulus field was accounted for when generating stimuli of a specified contrast.

Prior to data collection, the right eye of the participant was pharmacologically dilated with a 1% tropicamide ophthalmic solution following administration of 0.5% proparacaine as a local anesthetic. Subjects viewed the stimulus field through a 6 mm diameter artificial pupil.

### Psychophysical sensitivity task

Psychophysical sensitivity to flicker was measured using a two-interval forced-choice task. On each trial, one of two temporal intervals (the target interval) contained the target stimulus, which was a sinusoidal flicker (5 Hz) designed to stimulate the L, M and S cones. The other (reference) interval contained no modulation. Each interval was 500 ms in duration, and the start of each was indicated with an auditory cue. A 500 ms inter-stimulus interval identical to the reference interval was presented in between the stimulus intervals. Both the target and reference stimuli were refreshed at the same nominal frame rate, to minimize the possibility that an artifact related to stimulus updating would provide a cue to the target interval.

After the intervals, the participant indicated by keypad button press their judgment as to which interval contained the target. The response interval was untimed, and participants were allowed take a break or blink before responding; the next trial started immediately after each response. Participants did not receive feedback.

The contrast of the flicker stimulus was varied from trial to trial through three interleaved adaptive staircases. A correct response by the participant led to a decrease in nominal flicker contrast, while 3, 2, or 1 consecutive incorrect responses (depending on the staircase for that trial) led to an increase. The participant was asked to hold their eye in the eyepiece throughout a given experimental condition to maintain adaptation to the background.

During data collection, both the investigator (author JV) and the participant were masked to the order of the adaptation conditions in that session, the flicker contrast on any trial, and which interval contained the target stimulus. Particularly for those participants who were also investigators, it is possible that familiarity with the perceptual quality of the different conditions might have nonetheless allowed the participant to identify a particular condition.

### Threshold estimation

Pyschophysical sensitivity was assessed by measurement of the threshold contrast of a detectable flicker stimulus. Data for each condition for each session separately were fit with a cumulative Weibull psychometric function using the Palamedes 1.8.2 toolbox for MATLAB (Prins & Kingdom, 2018) for maximum likelihood fitting. These psychometric functions express the probability of detecting the flicker stimuli as a function of flicker contrast (Experiments 1 and 2), or of flicker stimulus RGB value (Experiment 3). The Weibull functions were fit to the data from all 120 trials in a block; guess rate was fixed at 50% for the two-interval forced-choice task, lapse rate was free to vary between 0-5%, and the slope parameter was unrestricted.

Threshold flicker contrast was estimated, separately for each condition for each session, from these psychometric functions, as the contrast corresponding to 70.71% correct detection. The 4 (or 6) repeated sessions (for Experiments 1 and 2, and for Experiment 3, respectively), yielded 4 (or 6) estimates of flicker detection threshold per participant for each melanopsin condition.

### Experiment 1

Experiment 1 measured the effect of adaptation to different levels of steady background melanopic content on sensitivity to cone-directed flicker. Flicker sensitivity was tested on a low-melanopic background and a high-melanopic background in separate blocks of trials, with each block starting with 5 minutes of adaptation to the background (Figure 2a, upper panel). As part of Experiment 1, similar measurements were carried out for low and high cone-directed backgrounds (with no change in melanopsin stimulation). This served as a positive control, demonstrating that the experimental manipulation leads to detectable Weber-like changes in sensitivity with background cone content (see Supplementary Information).

**Figure 2.**
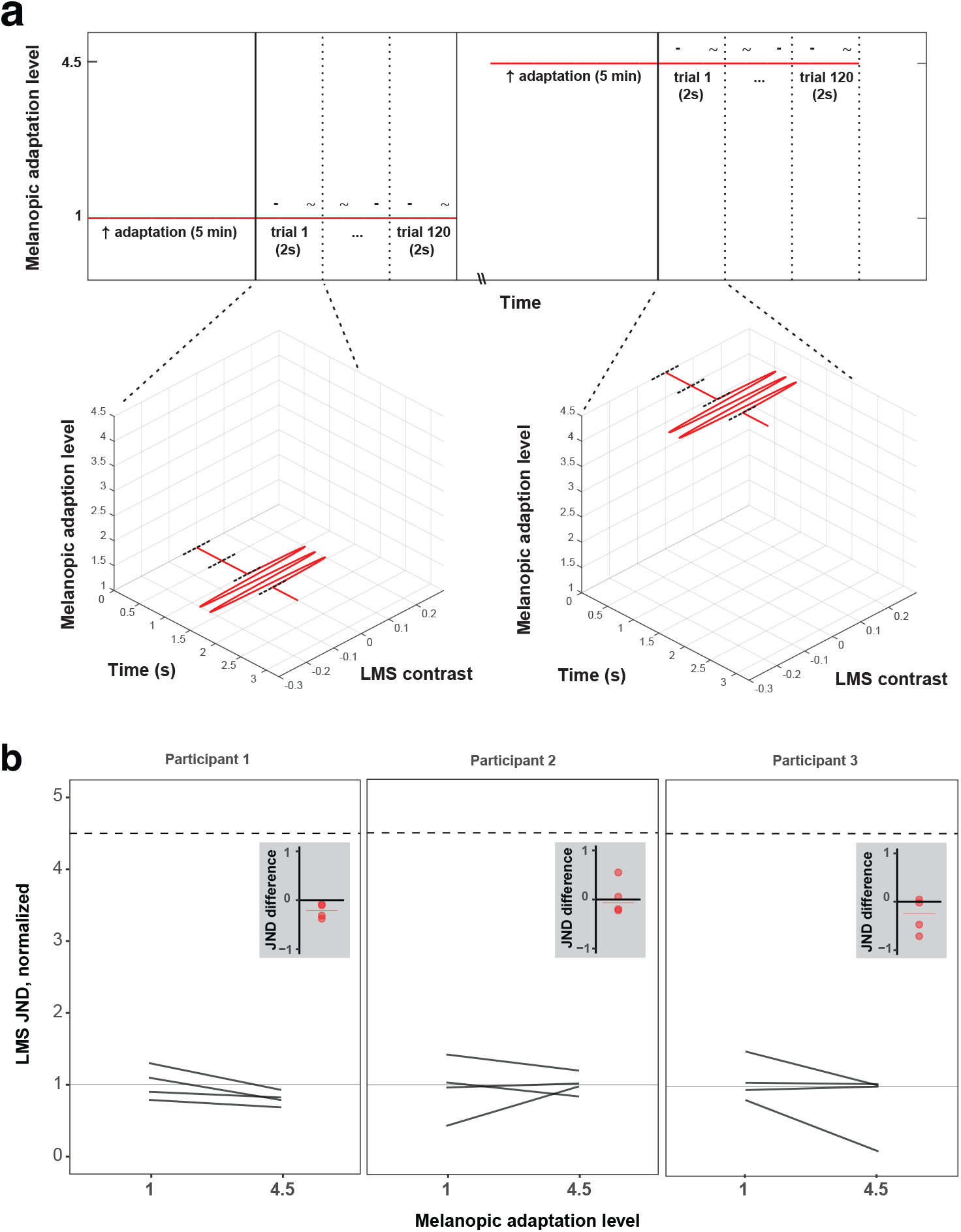
Experiment 1 stimulus and results. **a)** Each block contained only a single level of melanopic background adapting stimulus (vertical axis gives normalized level: 1 or 4.5) which remained constant through the entirety of that block. Blocks started with an adaptation period of 5 minutes, during which the participant fixated on the background for that block. Participants then completed 120 trials of a 2IFC task. Individual trials consisted of two 500 ms intervals, each followed by a 500ms ISI. During either the first or second interval, sinusoidal flicker (5 Hz) directed at the L, M, and S cones was presented around the melanopic background (low-melanopic background case shown on left, high-melanopic background case shown on right); during the other interval no such flicker was presented. After both intervals were presented, participants were asked to indicate which of the two intervals contained the flicker – this response was untimed. Intervals were indicated with an auditory cue. Participants could take a variable-length break in between blocks. Block-order was pseudorandom. **b)** Experiment 1 results. The plots show cone-directed flicker detection on low-and high-melanopic backgrounds, for the three participants in Experiment 1. Dark gray lines express detection as Just-Noticeable-Difference (JND) from the background in four single sessions. All JNDs for a participant were normalized to that participant’s median JND on the low-melanopic background. If detection of cone-directed flicker was mediated by melanopic stimulation in a Weber’s law fashion, JNDs on the 4.5x higher melanopic background would be 4.5x higher than on the low-melanopic background (dashed horizontal line). Insets: The four red dots show the within-session differences between JNDs at low-and high-melanopic adaptation level. The red horizontal line shows the median of the four within-session differences.

The target cone-directed flicker in this experiment was designed using silent substitution and delivered by the OneLight. For each background, a spectrum was designed to deliver 5% contrast in the positive direction (increase in excitation compared to background), and another to deliver 5% contrast in the negative direction (decrease in excitation compared to background), directed at the LMS cone photoreceptors (while keeping melanopsin excitation constant). The desired contrast of the cone-directed flicker on each trial was produced by scaling the two modulation spectra following the staircase procedure, providing a nominal contrast range of 0% to 5%, in 0.1% contrast steps. Sinusoidal variation in the relative weight of these components then produced flicker nominally directed at just the cone photoreceptors with the specified contrast. The background remained constant across all trials in a block (Figure 2a, lower panels).

Each session consisted of four blocks, each testing one background condition: the low-and high-melanopic backgrounds, as well as low and high cone-directed backgrounds that served to provide a positive control. The four blocks (one for each background) were acquired in pseudo-random order in each session. The participant adapted to each background for five minutes at the start of each block, and then completed 120 flicker-detection trials. Each participant completed 4 full valid sessions.

### Experiment 2

Experiment 2 measured the effect of a pulsed pedestal of melanopic content on sensitivity to cone-directed flicker. Flicker sensitivity was tested on interleaved trials on either a steady low-melanopic background, or on a pulsed high-melanopic pedestal (Figure 3a, upper panel). Each session started with 5 minutes of adaptation to the low-melanopic background, followed by 120 trials of each type (240 total) presented in pseudo-random order.

**Figure 3.**
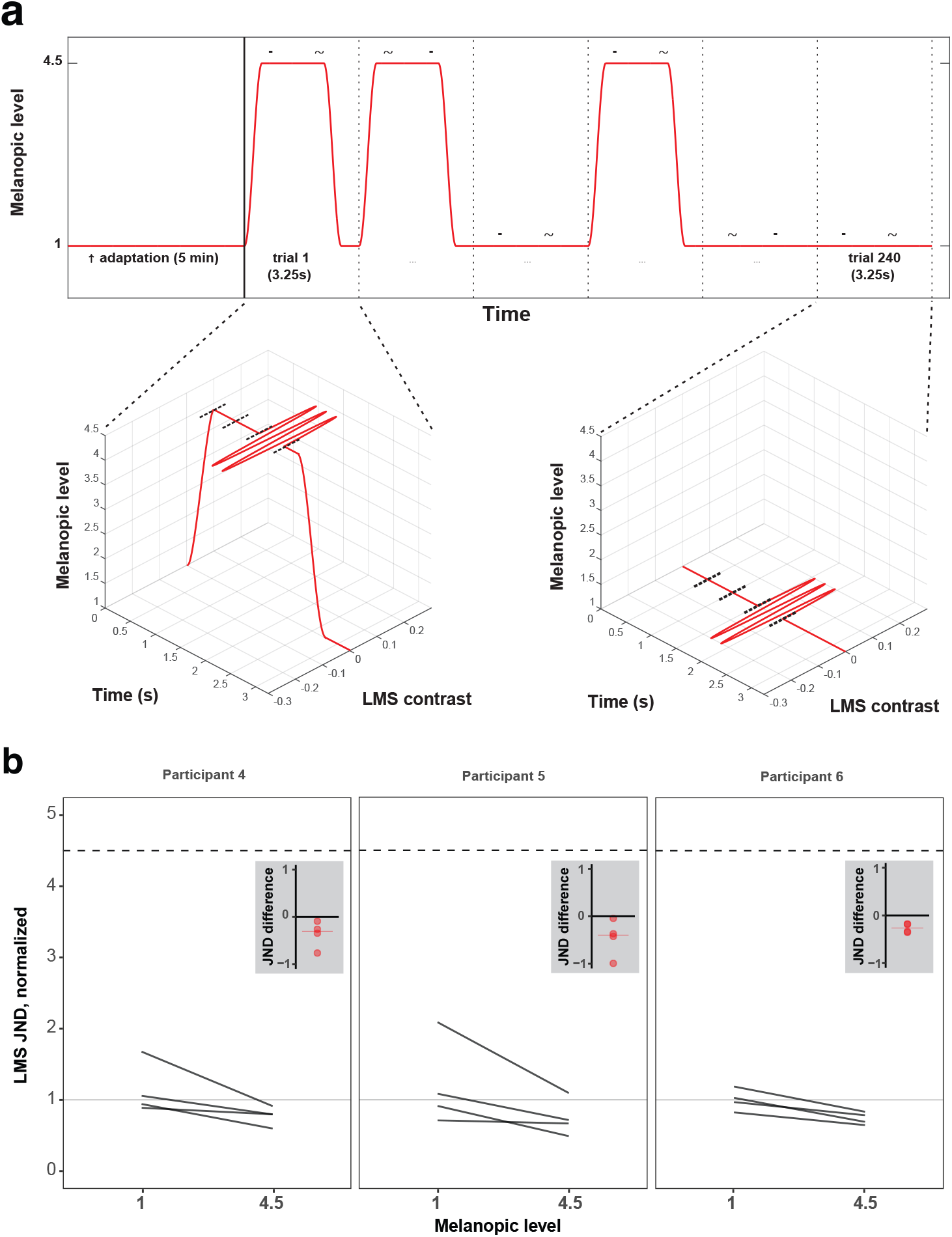
Experiment 2 stimulus and results. **a)** Temporal structure of trials in Experiments 2 and 3, where the target flicker was presented with or without a melanopsin-directed pedestal. Sessions started with an adaptation period of 5 minutes to the low-melanopic background. Participants then completed 240 trials of a 2IFC task, half of which contained a pedestal of melanopic stimulation. Individual trials consisted of two 500 ms intervals, separated by a 500 ms ISI. On trials containing the pedestal, melanopic stimulation rose during a 250 ms cosine window, and stayed high for the two intervals and ISI. During either the first or second interval, sinusoidal flicker directed at the L, M, and S cones was presented around the melanopic pedestal; on the other interval no such flicker was presented. After both intervals were presented, melanopic stimulation ramped back down to the low-melanopic background during another 250 ms cosine window, after which the participant was asked to indicate which of the two intervals contained the flicker – this response was untimed. On trials without the melanopic pedestal, melanopic stimulation stayed at the low-melanopic background level for the duration of the trial, including the 500 ms preceding the first interval. Intervals were indicated with an auditory cue. Participants could take a variable-length break in between blocks. Trial order was pseudorandom. **b)** Experiment 2 results. Same format as Figure 2c.

Half of the trials contained a pulsed melanopsin pedestal on which both stimulus intervals were presented (high-melanopic condition). The high-melanopic pedestal was windowed by a 500 ms half-cosine ramp. After 250 ms, the first interval was presented. After the second interval was presented, there was a 250 ms delay and then the melanopic content cosine-ramped back down to the low-melanopic background over the course of 500 ms. On the other half of trials the stimulus intervals were presented on a steady background with low-melanopic content (low-melanopic condition). On the trials without the melanopic pedestal, the steady background was presented for the combined 750 ms before the first interval, as well as for the combined 750 ms after the second interval (Figure 3a, lower panels).

The cone-directed flicker in this experiment was produced by the same method as in Experiment 1. Each session consisted of a single block of interleaved trials. Each participant completed 4 full, valid sessions.

### Experiment 3

Experiment 3 also measured the effect of a pulsed pedestal of melanopic stimulation on sensitivity to cone-directed flicker, but in Experiment 3 the target flicker was produced by the DLP projector optically admixed with the stimulus background/pedestal component produced by the OneLight. The target was thus a light-flux modulation rather than strictly cone-directed modulation. Flicker sensitivity was measured on either a steady low-melanopic background, or on a pulsed high-melanopic pedestal, on separate but interleaved trials (as in Experiment 2). Each block started with 5 minutes of adaptation to the low-melanopic background, followed by 120 trials of each type (240 total) presented in random order.

Melanopsin stimulation was produced in the same manner as in Experiments 1 and 2. The cone-directed light flicker was provided by the HP Notebook Companion digital light projector (DLP), under control of the same computer as the OneLight. The DLP was set at its half-on level (input settings = [0.5 0.5 0.5]) in the annular stimulus region, providing a DLP background component around which the flicker modulated. The 5 Hz sinusoidal flickering stimulus around this background was produced by time-varying the settings of the DLP both in the positive direction (increase in excitation compared to background) and the negative direction (decrease in excitation compared to background). The maximum excursion, i.e., the change in input settings, was adjusted between trials.

The primary interest in Experiment 3 was to confirm the results of Experiment 2, but with a different stimulus generation method. In this stimulus generation method, the same flicker stimuli from the projector were admixed across the introduction of the melanopic pedestal. This did not require precise control over the projector, as the goal was simply to make the test stimulus independent of the background/pedestal. In particular, no linearization of the DLP stimulus was performed. Rather, the RGB input to the projector was varied at a nominal 8-bit quantization, with the R, G and B inputs always set to the same value as each other. Additional quantization may have been performed internally to the DLP. Flicker stimuli were drawn at a 60 Hz refresh rate.

As described above, the output intensity of the DLP was reduced by passing it through 3.3-log of neutral density filter, then imaged onto a custom diffuser which acted as a projection screen, and finally admixed to the optical signal from the OneLight by passing it through the beam splitter. The participant saw the full 27.5 degree annulus stimulus region as a mix of the OneLight stimulus component, and the DLP stimulus component, with the central 5 degrees and outer region masked by a dim white light from the DLP.

The dynamics of the melanopsin stimulation and the trial design were the same as in Experiment 2 (see Figure 3a). Each session consisted of a single block of interleaved trials (120 trials with and without pedestal, 240 total). Each participant completed 6 validated sessions.

## Results

In each of three experiments, sensitivity to cone-directed flicker was measured in low and high melanopsin stimulation conditions. In Experiment 1, melanopsin stimulation was varied by adaptation to either a low-or high-melanopic background, and the detection task was completed for each condition separately. In Experiments 2 and 3, melanopsin stimulation was varied on individual, intermixed trials by presenting the cone-directed flicker around a low-melanopic background, or upon a high-melanopic pedestal. In Experiments 1 and 2, a single device (the OneLight) was used to both modulate melanopic excitation and to produce cone-directed flicker. In Experiment 3, the target flicker was produced by a conventional, 3 primary projector, and optically admixed with the background/pedestal produced by the OneLight.

In Experiment 1 and 2, threshold was measured as the estimated contrast for 70.71% correct detection of the flicker stimulus, separately for each melanopsin condition of each session. Table 1 presents the LMS threshold contrasts in each melanopsin condition, expressed as the median across sessions per participant.

**Table 1:**
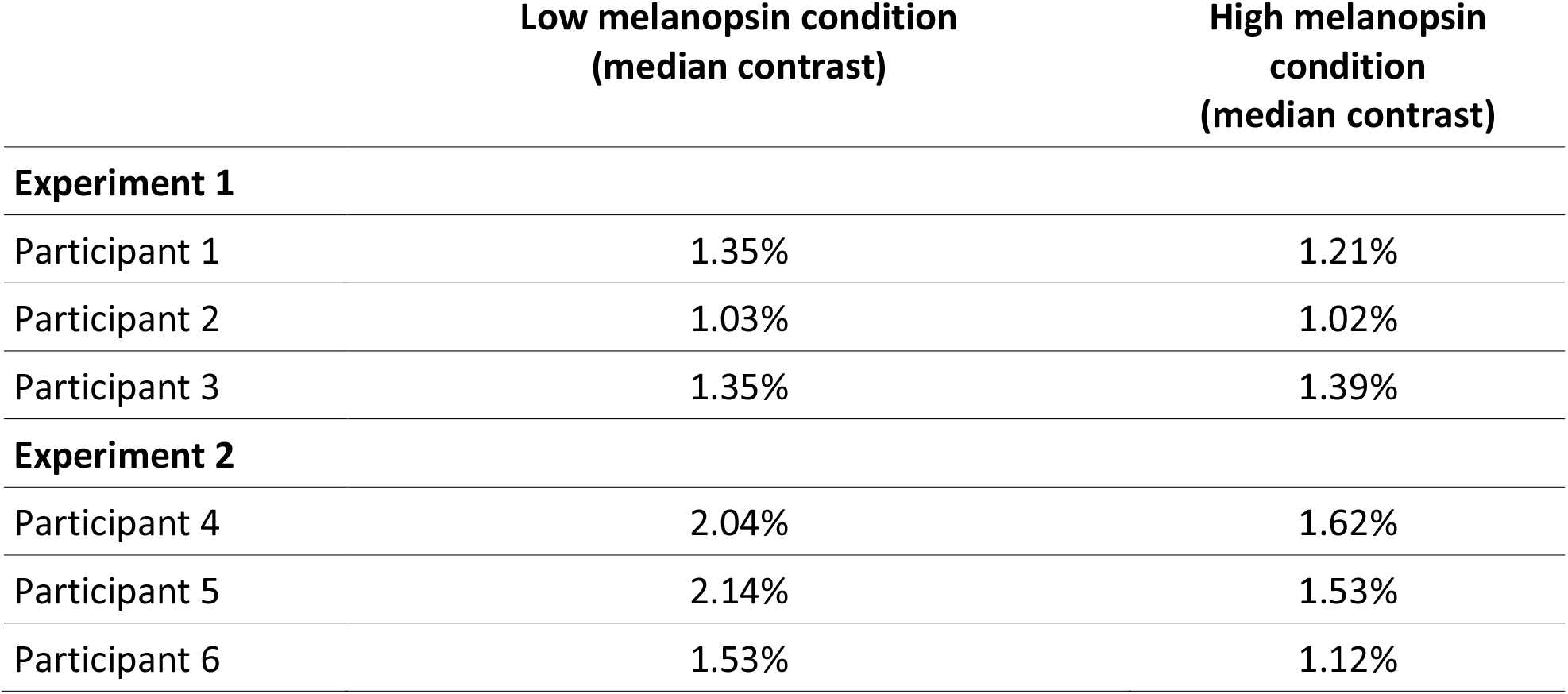
participant flicker LMS contrasts at detection threshold. Median (over sessions) threshold contrast for LMS flicker in low, and high melanopsin conditions.

|  | Low melanopsin condition<br>(median contrast) | High melanopsin<br>condition<br>(median contrast) |
| --- | --- | --- |
| **Experiment 1** |  |  |
| Participant 1 | 1.35% | 1.21% |
| Participant 2 | 1.03% | 1.02% |
| Participant 3 | 1.35% | 1.39% |
| **Experiment 2** |  |  |
| Participant 4 | 2.04% | 1.62% |
| Participant 5 | 2.14% | 1.53% |
| Participant 6 | 1.53% | 1.12% |

Additionally, sensitivity can be expressed as the Just-Noticeable-Difference (JNDs) between flicker stimulus and background. Multiplying contrast values by the LMS content of the background spectrum gives the increase and decrease in LMS stimulation produced by the flicker stimulus. Thus, multiplying threshold contrast values by background LMS content expresses detection threshold as the difference in LMS stimulation between background and stimulus that is just noticeable.

In visualizations of results (Figures 2b & 3b), JNDs are normalized such that the median JND in the low melanopsin stimulation condition has a value of unity. This allows for the interpretation of the JNDs in the high melanopsin condition as a scalar increase of JNDs in the low melanopsin condition. The change in melanopic contrast was 350%; thus melanopsin stimulation was increased by a factor of 4.5x between the low-and high-melanopic backgrounds. If, for example, background melanopsin contrast had the same effect upon LMS sensitivity as background luminance contrast itself, then the JNDs in the high-melanopic condition would be expected to be 4.5, following Weber’s Law. Central tendency is visualized as the median JNDs for each condition.

In the current experiments, the effect of melanopsin on detection of cone-directed signals was investigated within participants, by varying the melanopsin stimulation within each session. Thus, to determine the presence or absence of an effect of the melanopsin stimulation, the difference between JNDs in the high and low melanopsin condition was calculated for each session (Figures 2b & 3b, insets). If melanopsin has no effect on detection, the JNDs would not differ systematically between melanopic conditions, and these within-session differences would cluster around 0.

### Experiment 1

In Experiment 1, the threshold for two-interval discrimination of cone-directed flicker was measured on a low-or a high-melanopic background (Figure 2a). participants adapted to the low-or high-melanopic background for five minutes, and the background was held constant during collection of 120 trials.

Detection thresholds for cone-directed flicker on the constant high-melanopic background were not systematically different from detection thresholds on the constant low-melanopic background (Figure 2b). For participants 1-3, median JND in the high melanopsin stimulation condition were, respectively, 80%, 100%, and 100% percent of the median JND in the low melanopsin stimulation conditions. For each participant, the difference in JNDs between melanopsin conditions is small compared to the variation in these differences across sessions (Figure 2b, insets). Therefore, no reliable difference in JNDs of cone-directed flicker between the low and high melanopsin conditions was found; one participant appears to show a median decrease in sensitivity, while the other two participants show no systematic difference in median threshold.

The lack of change in thresholds was not due to lack of power in the procedures used to measure such effects. Control data were collected as part of Experiment 1, in which the L, M, and S content of the backgrounds differed but their melanopic content was held constant. In this control condition, thresholds for the LMS-directed flicker increased as expected from Weber’s Law (see Supplementary Information).

### Experiment 2

In Experiment 2, the threshold for two-interval discrimination of cone-directed flicker was measured on a low-melanopic background, or a pulsed high-melanopic pedestal (Figure 3a). Participants adapted to the low-melanopic background for five minutes, and on individual trials were presented either with only the low-melanopic background or the high-melanopic pulsed pedestal on top of that background.

Figure 3b presents the thresholds for cone-directed flicker on a pulsed high-melanopic pedestal as compared to the low-melanopic background. For participants 4-6, median JND in the high melanopsin stimulation condition were, respectively, 80%, 69%, and 74% percent of the median JNDs in the low melanopsin stimulation conditions For all three participants, there was a small but consistent tendency for JNDs to be reduced on the high as compared to low melanopic background. The within-participant differences in JNDs between melanopsin conditions are shown in the insets (Figure 3b).

Experiment 2 seems to show a small, consistent decrease in JNDs of cone-directed flicker between the high and low melanopsin conditions. However, it could not be ruled out that this difference between conditions was due to confounding stimulus artefacts (rather than a true effect of melanopsin stimulation).

### Potentially confounding stimulus artefacts

The results of Experiments 1 and 2 show either no effect, or a small effect, of melanopsin background upon cone flicker detection. Two alternative hypotheses suggest that either there is no true effect of melanopsin stimulation (as evidenced in experiment 1), and the apparent effect in experiment 2 is due to confounds; or, alternatively, experiment 2 shows a true effect that is masked by confounds in experiment 1. The measurement could be confounded in such ways if stimuli depart from their nominal spectral content. There could be imprecision in the LMS content of low-and high-melanopic backgrounds or in the magnitude of LMS flicker contrast itself, either of which could cause the LMS flicker to have contrast different from that intended on the two backgrounds. Further, stimulus imprecision could result in the stimulus being detected by a mechanism other than the one targeted (e.g., the highly sensitive L–M cone opponent mechanism). Spectroradiometric measurements of the stimuli were obtained during each session of data collection, and these measurements were used to assess each of these potential confounds.

### Imprecision in background spectra

In Experiments 1 and 2, the low-melanopic and high-melanopic spectra were designed through silent substitution that targeted melanopsin while attempting to silence the L, M and S cones. Imprecision in device control affects the efficacy of this silencing: residual contrast (“splatter”) on the L, M, and S cones could have been present across the nominally cone-silent background exchange. Since the pair of spectra was generated separately for each session of data collection, the imprecisions varied session-to-session. Radiometric measurements of the background spectra were taken at the end of each session of data collection to assess the magnitude of the background splatter.

Figure 4a, left, shows the measured LMS content of the low-melanopic and high-melanopic backgrounds in Experiment 1. LMS content here is the sum of L, M and S-cone excitations. Excitations were calculated by multiplying measured spectral power distributions by cone fundamentals (normalized such that they peak at 1). For all three participants, the LMS content does not vary systematically between the melanopic backgrounds.

**Figure 4.**
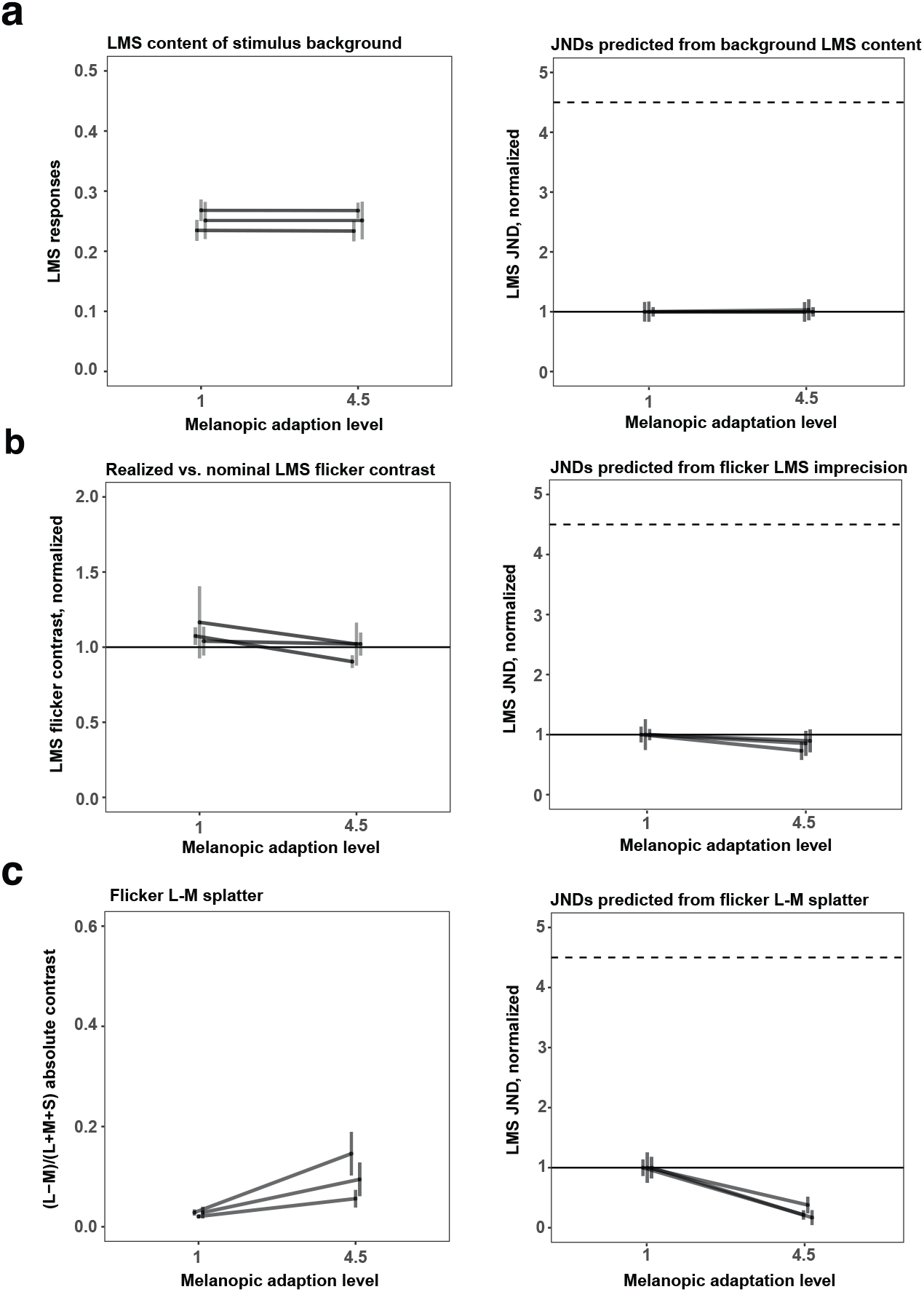
Stimulus imprecision analysis for Experiment 1. **a)** Left panel. LMS content of the low-melanopic and high-melanopic backgrounds in Experiment 1, in cone coordinates. Stimulus spectra were multiplied by cone fundamentals (normalized to peak at 1) to obtain coordinates for each cone class; these were then averaged to get a combined LMS cone coordinate. Right panel. Predicted JNDs if background LMS content were driving LMS detection threshold, under a Weber’s Law assumption for detection (JND proportional to background LMS content). Same format as Figure 2c, but with median (across sessions) JNDs and +/-1 standard error of the median for each of three participants shown in a single plot. **b)** Measured LMS content of the threshold flicker stimuli in Experiment 1. Left panel. Measured LMS content of the flicker stimuli corresponding to detection threshold in the low-melanopic and high-melanopic conditions, expressed as proportion of the nominal LMS contrast of those flicker spectra (i.e., value of 1 indicates LMS contrast was as nominal). Each line plots median +/-1 standard error of the median for one participant, with the median taken across sessions. Right panel. JNDs for Experiment 1, based on measured rather than nominal flicker spectra. **c)** L-M chromatic contrast of the LMS flicker stimulus spectra in Experiment 1. Left panel. Measured (L-M)/(LMS) contrast of the flicker stimuli in the low-melanopic and high-melanopic conditions. Each line plots median +/- 1 standard error of the median for one participant, with the median taken across sessions. Right panel. Predicted JNDs, assuming that detection of the flicker stimulus was mediated not by LMS contrast, but rather by the absolute magnitude of the L-M chromatic contrast. Under this assumption, greater L-M contrast in the flicker stimulus predicts lower JNDs, under the null-hypothesis that cone-mediated flicker detection does not change with background melanopsin stimulation.

To understand the effect of any difference in cone stimulation between backgrounds on the psychophysical sensitivity, normalized JNDs were predicted for each session. This prediction was made as follows: First, a difference in melanopsin stimulation was assumed to have no effect on sensitivity. The nominal LMS contrast at threshold, as measured on the low-melanopic background, was taken as the predicted threshold contrast for both backgrounds, i.e., the threshold flicker LMS contrast was assumed constant across melanopic conditions. Second, this predicted contrast was multiplied with the measured LMS content of each background. This expresses the predicted threshold as the predicted JND on each background. The predicted JNDs were normalized in the same way as the experimental results, such that the median (per participant) JND in the low melanopsin stimulation condition has a value of 1. These normalized predicted JNDs express thresholds as the increase in LMS content that would correspond to the same predicted LMS contrast on each background, given the measurements of that background, under a Weber’s law assumption.

Figure 4a, right, shows the median (per participant) of these normalized, predicted JNDs. They do not change appreciably across the two melanopic backgrounds, confirming that the measured LMS contents of the low-and high-melanopic backgrounds do not differ enough to lead to substantially different JNDs. Thus, imprecisions in the nominally constant LMS content of the backgrounds would not produce an effect that could be masking a true effect of melanopsin stimulation on psychophysical sensitivity to cone-directed flicker and leading to the observed null-result.

Figure 5a shows the same analysis, with the same conclusion of no artifact, for Experiment 2.

**Figure 5.**
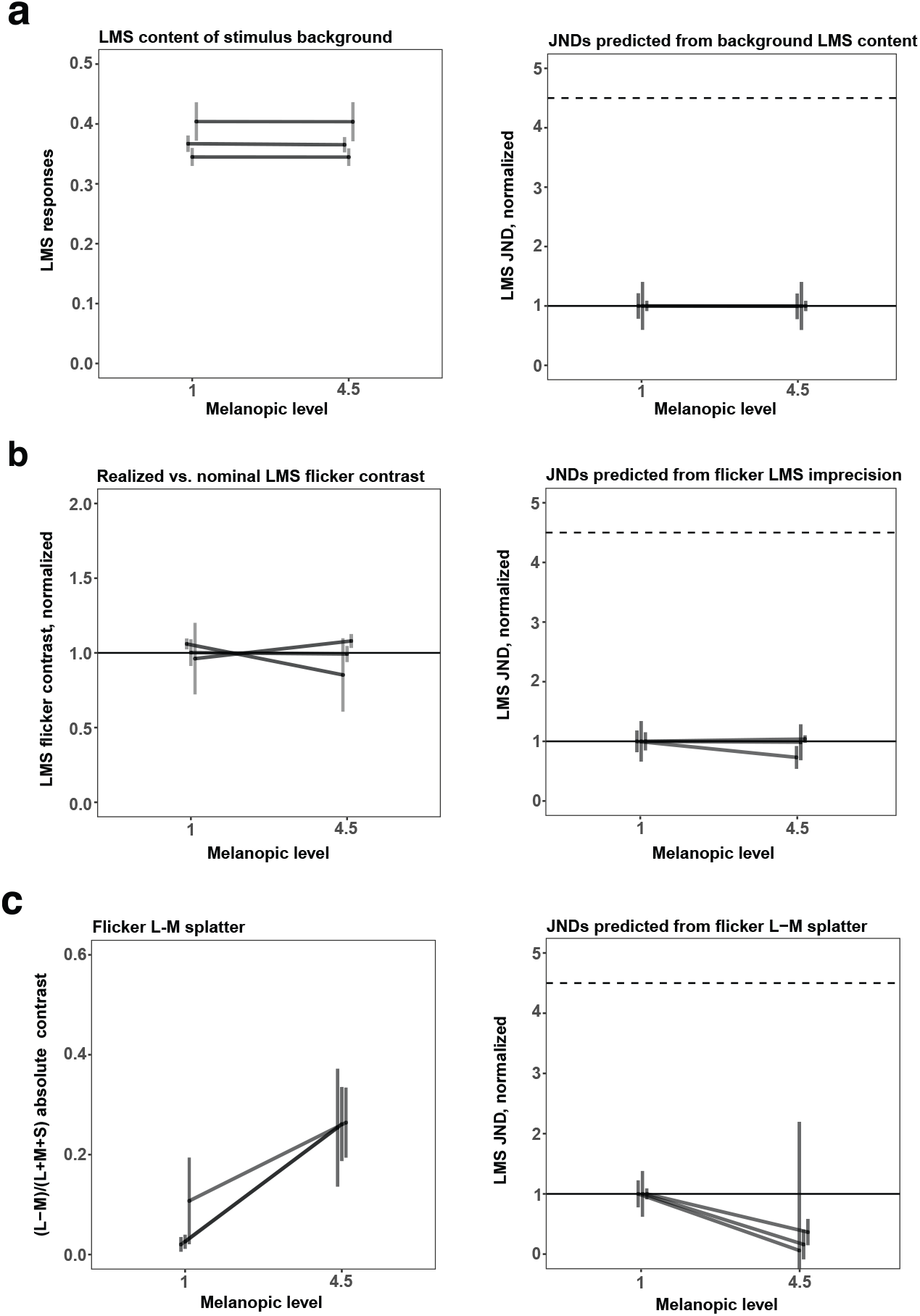
Stimulus imprecision analysis for Experiment 2. Same format as Figure 4.

### Imprecision in flicker

The flicker stimuli in Experiments 1 and 2 were also designed through silent substitution to produce equivalent modulation of the L, M and S cones on both melanopic backgrounds. Radiometric measurements of the flicker spectra were also taken, at the end of each session. A preliminary nominal threshold was estimated from the staircase reversals – not from a psychometric function fit – and this preliminary nominal threshold was used to generate stimulus spectra corresponding to the nominal stimulus at detection threshold. From these spectra, LMS flicker contrast was calculated as the equally weighted sum of L, M, and S-cone contrasts between these spectra and the corresponding measured background spectra.

Figure 4b, left, shows the measured flicker LMS contrasts, in the low-and high-melanopic conditions. Here the measured contrast is expressed as a proportion of the nominal contrast. In the ideal case of no splatter, this proportion would equal 1 (measured flicker LMS contrast equals nominal flicker LMS contrast); higher values indicate the flicker stimulus had more cone-directed content than nominal, lower values indicate less than nominal content. On average, the flicker was slightly stronger than nominal in the low-melanopic condition, but closer to nominal in the high-melanopic condition, although this is not consistent or robust across participants and sessions. This suggests that the same nominal flicker threshold corresponds to a weaker flicker stimulus in the high-melanopic condition than in the low-melanopic condition, thus potentially overestimating the sensitivity in the high-melanopic condition.

To understand the effect of this increase on the psychophysical sensitivity, JNDs were predicted for each session. This prediction was made as follows. First, a difference in melanopsin stimulation was assumed to have no effect on sensitivity. The nominal LMS contrast at threshold, as measured on the low-melanopic background, was taken as the predicted threshold contrast for both backgrounds, i.e., the threshold flicker LMS contrast was assumed constant across melanopic conditions. Second, this predicted contrast was multiplied with the ratio between measured and nominal LMS contrast of each flicker stimulus. This, then, predicts the LMS contrast (including splatter) of the flicker stimulus, at the predicted threshold. Lastly, this predicted contrast was multiplied with the measured LMS content of each background. This converts the predicted threshold into the predicted JND on each background.

Figure 4b, right, shows these predicted JNDs, averaged per participant as the median over sessions for Experiment 1. These differ minimally across the two melanopic backgrounds, indicating that imprecisions in the nominally identical flicker stimuli did not produce an effect that could be masking a true effect of melanopsin stimulation on psychophysical sensitivity to cone-directed flicker and thus producing an artifactual null-result. If anything, the splatter analysis indicates that a true lack of effect might be revealed as a small decrease in JND with increasing background melanopic content, consistent with the weak trend seen in the data for Experiment 1. Figure 5b shows the same analysis, with similar conclusion, for Experiment 2.

Imprecise production of the LMS flicker stimuli could also have introduced L-M chromatic contrast, to which the visual system is highly sensitive (Chaparro et al., 1993). Given enough L-M splatter, detection of the flicker stimulus could have been mediated primarily by this chromatic contrast, rather than by the experimentally manipulated LMS contrast. From the same radiometric measurements of the flicker stimuli, the L-M contrast splatter was computed. To compute the L-M contrast for other nominal values of L+M+S stimulus, we express the L-M splatter relative to the measured L+M+S contrast, as the contrast ratio (L-M)/(L+M+S). We scale this ratio by the nominal value of L+M+S to obtain an estimate of L-M splatter as nominal target contrast is varied.

There was a trend for the (L-M)/(L+M+S) ratio of the flicker contrasts to be lower for the low-melanopic condition than for the high-melanopic condition, for all participants in both Experiment 1 (Figure 4C, left) and Experiment 2 (Figure 5C, left). To understand the effect of this chromatic contrast on the psychophysical sensitivity, JNDs were predicted for each session, under the null-hypothesis that melanopic content does not affect sensitivity and that detection of the flicker stimulus was mediated exclusively by the L-M chromatic contrast.

Specifically, for each session the absolute value of measured L-M contrast at threshold on the low-melanopic background was taken as the predicted threshold contrast for both conditions, i.e., the L-M contrast was assumed constant at threshold across melanopic conditions. By multiplying this predicted contrast with the ratio between L-M and LMS contrast, the nominal LMS contrast of a flicker stimulus with that predicted L-M contrast on the high-melanopic background was calculated. If the L-M splatter mediated detection and there were no effect of background melanopsin content on thresholds, a decrease in threshold (increase in sensitivity) would be observed on the high-melanopic background. Figure 5c, right shows the same analysis, with the same conclusion, for Experiment 2.

The results from Experiment 2 potentially reflect this kind of stimulus artifact. Overall, participants showed slightly increased sensitivity for the flickering stimulus on the high melanopsin background. While this could in principle be an effect of melanopsin excitation, such an effect in the data is consistent with the direction of effects produced by the small amount of splatter in the stimuli, so that the small changes in threshold sensitivity might be artifactual. To address this possibility, a third experiment was conducted, in which the generation of the stimulus flicker was independent of the manipulation of the stimulus background.

### Experiment 3

Experiment 3 was designed to address the possibility that imprecision in the flicker stimulus could confound the perceptual measurement on different melanopic backgrounds. For this experiment, the low-melanopic background and high-melanopic pedestal were produced by silent-substitution using the OneLight as in Experiment 2. The flicker stimuli, however, were generated using the DLP projector, a device separate from that used to produce the background and pedestal, and optically admixed. Thus, the spectral content of the flicker stimuli was held constant across melanopic conditions (session-to-session variation may have occurred, but this variation would be independent of, and not correlated with, the melanopic condition).

In Experiment 3 the threshold for two-interval discrimination of cone-directed flicker was measured on a low-melanopic background and on a pulsed high-melanopic pedestal (as in Experiment 2). Participants adapted to the low-melanopic background for five minutes, and on individual trials were presented either with only the low-melanopic background or the high-melanopic pulsed pedestal on top of that background.

Detection thresholds for cone-directed flicker admixed on a pulsed high-melanopic pedestal, were not systematically different than detection thresholds on a low-melanopic background For participants P1, P4 and P5, median JNDs in the high melanopsin stimulation condition were, respectively, 106%, 100% and 89% of the JNDs in the low melanopsin stimulation conditions (Figure 6). For each participant, the difference in JNDs between melanopsin conditions is small compared to the variation in these differences across sessions (Figure 6, insets). Therefore, no reliable difference in JNDs of cone-directed flicker between the low and high melanopsin conditions was found.

**Figure 6.**
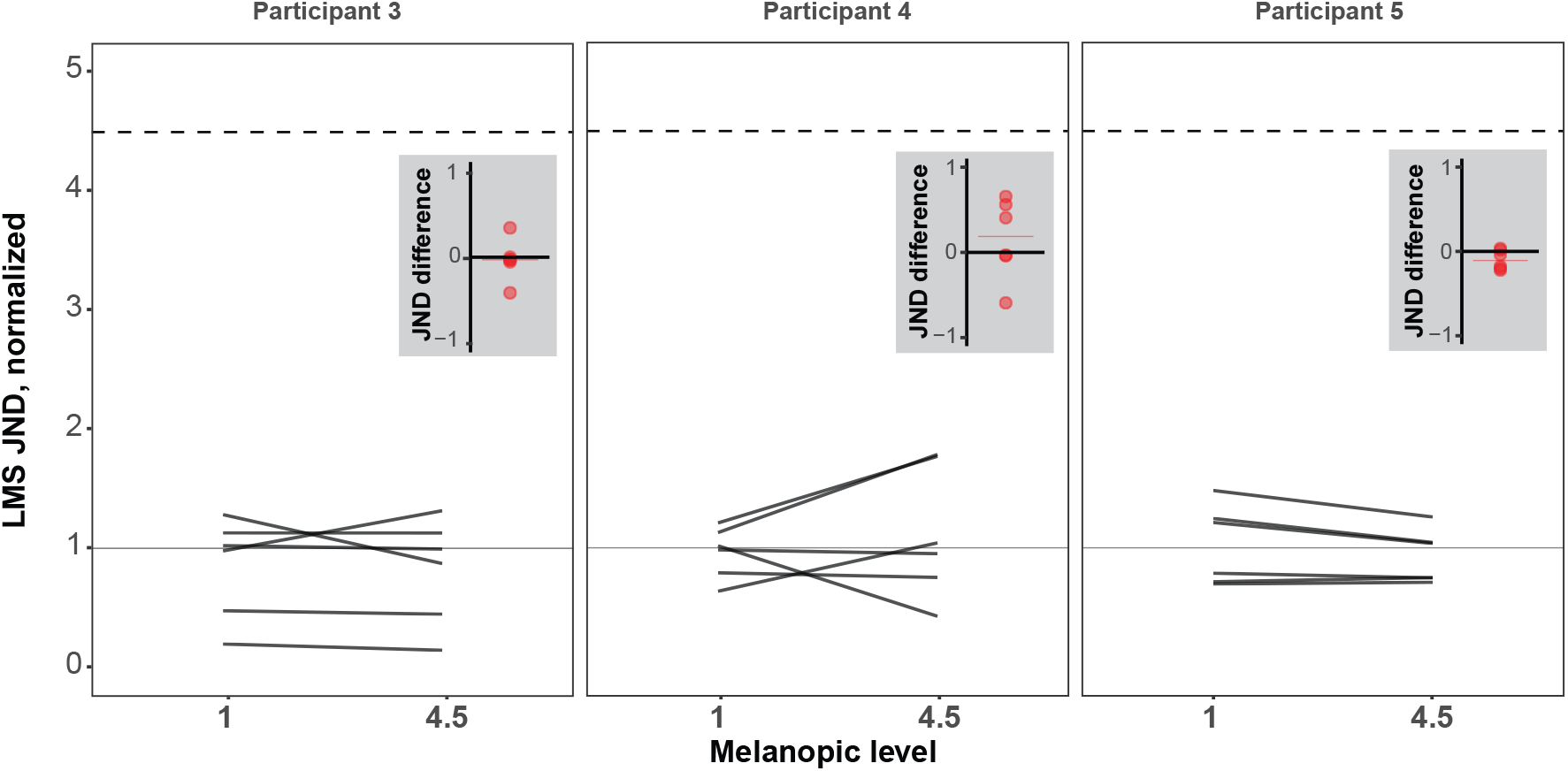
Experiment 3 results. Same format as Figure 2c.

## Discussion

We find that varying melanopsin stimulation does not measurably alter threshold sensitivity for cone-directed luminance flicker. Interaction effects are absent both during adaptation to a continuous, high melanopsin background (Experiment 1), and against brief, high melanopsin spectral pulses (Experiments 2 and 3). This null result was obtained in the face of slight imperfections in our control of the chromatic content of our stimuli. We consider it notable that failure to measure and account for these small artifacts might have otherwise led to the erroneous conclusion of an effect of melanopsin stimulation upon cone sensitivity.

Melanopsin resides within the ipRGCs, and these cells are known to project to the geniculo-striate visual pathway (Dacey et al., 2005). There is evidence that melanopsin stimulation contributes to perceived brightness, in addition to the perception of brightness provided by the cone-mediated luminance mechanism (Brown, Tsujimura et al. 2012; Spitschan et al. 2017; Zele et al. 2018a; Yamakawa et al, 2019; DeLawyer et al., 2020). This brightness percept follows the properties predicted from ipRGC physiology, including stronger percepts at higher background light levels and lower temporal frequencies (Lucas et al., 2020). A related finding is that cone and melanopsin signals combine in a sub-additive manner in producing ratings of visual discomfort (McAdams et al., 2020). These photoreceptor signals must be combined somewhere prior to sites mediating verbal report, and in principle this combination could begin within the ipRGCs themselves, as several classes of these cells receive extrinsic cone inputs.

We sought to test in the current study the specific proposition that melanopic signals from the ipRGCs interact in human perception with cone-driven signals carried by the classic RGCs. Specifically, we began with the assumption that cone-directed luminance flicker is processed predominantly by the classic RGCs. If manipulation of the melanopic background (which can only be transduced by the ipRGCs) alters perceptual sensitivity for luminance flicker, this would provide support for the idea that ipRGC and classic RGC signals interact in perception (although with some caveats discussed below). Across our three experiments, we did not find reliable evidence of a change in threshold cone sensitivity induced by changing melanopsin activation. While it is of course possible that a small interaction effect exists that escaped the sensitivity of our measurements, our findings nonetheless place tight bounds on how much interaction there could be between melanopsin stimulation and luminance flicker detection at threshold.

An inferential nuance of our study is that a positive finding in our measurements would not have provided definitive evidence of an interaction specifically of ipRGC signals with those of the classic RGCs. First, it is possible that the 5 Hz luminance modulation could be transduced by the ipRGCs themselves. Second, because Experiment 3 made use of a light-flux flicker modulation (as opposed to pure luminance) it is possible that the melanopic component of the light-flux flicker could have directly modulated ipRGC activity. Both of these considerations admit the possibility that the background and flicker components of the stimuli could have interacted entirely within the ipRGCs. In practice, we think that both of these theoretical mechanisms are unlikely to contribute much to perception on prior grounds, and more generally these considerations are mooted by our negative results.

Prior studies with animals and humans have suggested that ipRGC and classic RGC signals can interact. In rodents, a ten-fold increase in melanopic background leads to a measurable decline in the b-wave (post-photoreceptor) ERG response to a brief flash of light (Allen et al., 2014); a similar effect of melanopsin excitation on global mean activity in the LGN was observed. This effect is proposed to arise from recurrent axon collaterals from the ipRGCs that modulate cone inputs to the bipolar cells (Joo et al., 2013). There are salient differences between our perceptual study in humans and this electrophysiologic measurement in rodents that may explain the different findings of an effect of melanopsin background. While our study provided substantial (350%) contrast on melanopsin between the low and high background conditions, our stimulus manipulation is nonetheless unable to achieve the higher contrast possible in transgenic mice. It is possible that an effect of melanopsin background upon cone sensitivity arises only for changes in background beyond what we were able to achieve. Additionally, we examined perception of a subtle modulation of cone signals at perceptual threshold, as opposed to the supra-threshold, high-intensity flash of light used in the ERG measurement. It is possible that the effect of melanopsin activity upon cone signals becomes evident only in the supra-threshold case. We also cannot discount the possibility that interactions between melanopsin and cone signals are fundamentally different in humans than they are in the rodent model system.

The prior study most similar to our own is that of Zele and colleagues (2019). They measured human psychophysical sensitivity to one second, cone-directed pulses, either in the presence or absence of a melanopsin pedestal. In different measurements, contrast on melanopsin was varied between 9% and 22%. Despite this relatively small degree of melanopsin contrast, Zele and colleagues reported that the high melanopic pedestal induced a 40-60% increase in sensitivity to cone-directed stimuli. It is difficult to reconcile these findings with the negative results of the current study. While there were differences in the cone-directed stimuli (steady pulses vs. 5 Hz flicker) and background light level (637 cd/m2 in Zele and colleagues vs. ∼275 cd/m2 in the current study), these do not strike us as fundamental differences for a posited interaction of ipRGC and classic RGC signals. In particular, these small stimulus differences seem unlikely to explain why our study found no effect of a large manipulation of melanopsin contrast upon cone-mediated perception, while Zele and colleagues found a large effect using a small change in melanopsin contrast.

We were aware in the current study of the particular importance of measuring and accounting for even small imprecision in the generation of stimuli. The current experiments (similar to the studies of Allen et al., 2014 and Zele et al., 2019) make use of the technique of silent substitution (Estévez & Spekreijse, 1982) to selectively target photoreceptor classes. As we have described in this report, inevitable imperfections in device control produce deviations of the stimuli from their nominal properties. Most critically, we found that our nominal luminance isolating flicker stimulus in Experiments 1 and 2 developed ∼0.5% of L-M chromatic contrast “splatter” when it was generated on the high melanopsin background spectrum. While a seemingly small degree of stimulus error, we calculated that this nonetheless could lead subjects to exhibit increased sensitivity for detecting the flicker on the high melanopsin background. This observation motivated Experiment 3, in which the design of the apparatus precluded an influence of the melanopsin background upon the flicker stimulus. This third experiment allowed us to reject the presence of an interaction in perception of pulses of melanopsin stimulation and threshold sensitivity for cone-directed flicker.

We consider it possible that the results reported by Zele and colleagues (2019) contain an influence of stimulus imprecision of this kind, although we note that the authors did consider and reject this possibility. A similar note of caution should likely attend claims that melanopsin stimulation imparts or alters chromatic percepts (Cao et al. 2018; Zele et al. 2018b; see Brown et al. 2012 and Spitschan et al., 2017).

In summary, we did not find in the current study evidence that melanopsin-driven signals arising from the ipRGCs influence human perception of cone signals conveyed by the classical RGCs. It remains possible that such interactions exist for stimulus regimes different from those we examined. In particular, prior work in rodents leads us to expect that, if such interactions are to be found, they are present for supra-threshold cone stimulation against very high melanopic backgrounds.

## Acknowledgements

This work was supported by grants from the National Eye Institute (R01EY024681 to GKA and DHB; Core Grant for Vision Research P30 EY001583) and the Department of Defense (W81XWH-151-0447 to GKA).

## Author Contributions

JV, DB and GKA designed the experiments, and participated in some of the experiments; corresponding author JV implemented the experimental design in software, collected data, analyzed data, and drafted the methods and results; Joint supervisors DB and GKA drafted discussion and introduction, and edited methods, results; EBH designed figures and provided copy-editing. All authors reviewed the manuscript.

## Data availability

These experiments were the subject of pre-registration documents (E1: https://osf.io/jd4qh/ ; E2: https://osf.io/qgs49/ ; E3: https://osf.io/p45aj/). The datasets and code used for data analysis is publicly available (https://github.com/BrainardLab/MeLMSens).

