## Supplemental Figure 1 for "Melanopic stimulation does not affect psychophysical threshold sensitivity for luminance flicker"

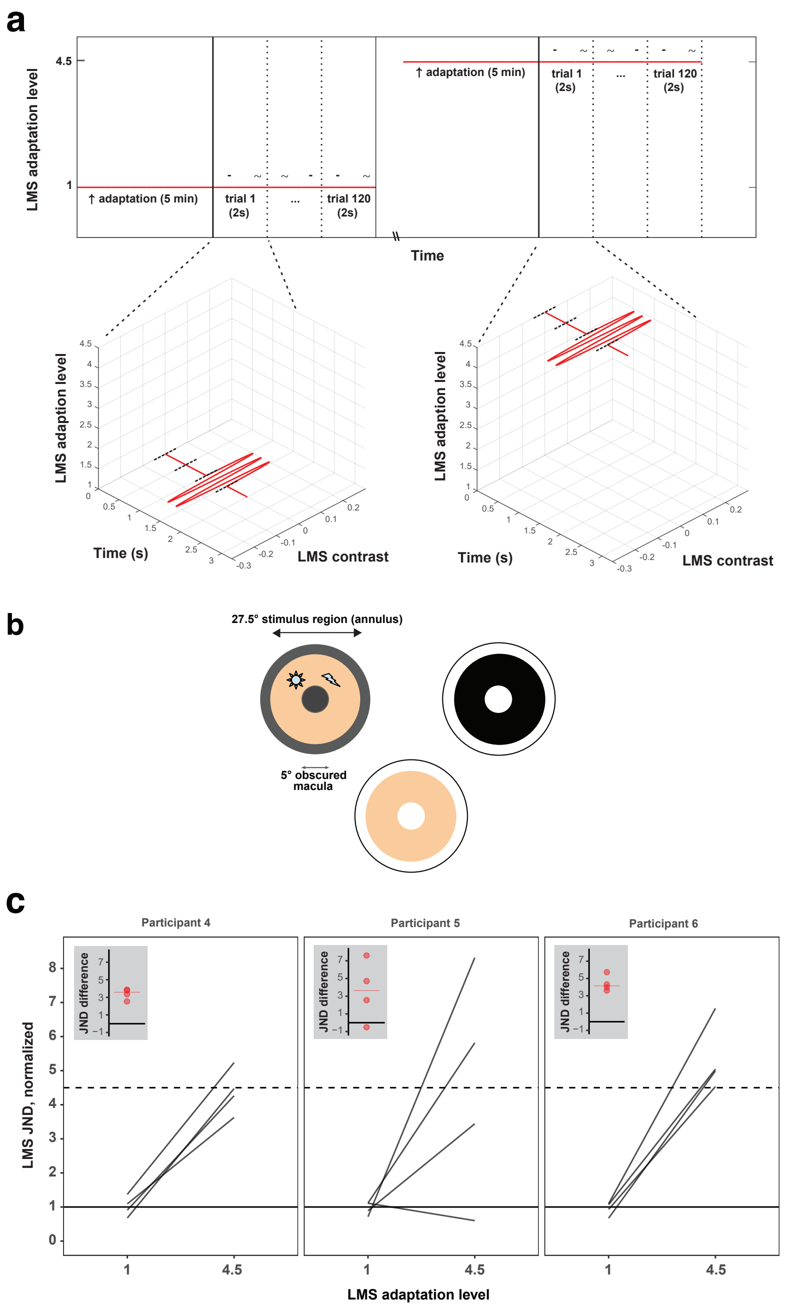


**Figure S1. Experiment 1 positive control stimulus and results**. **a)** Each block contained only a single level of L+M+S-cone directed stimulation (vertical axis gives normalized luminance: 1 or 4.5) which remained constant through the entirety of that block. Blocks started with an adaptation period of 5 minutes, during which the participant fixated on the background for that block. Participants then completed 120 trials of a 2IFC task. Individual trials consisted of two 500 ms intervals, each followed by a 500 ms ISI. During either the first or second interval, sinusoidal flicker (5 Hz) directed at the L, M, and S cones was presented around the background (low-LMS background case shown on left, high-LMS background case shown on right); during the other interval no such flicker was presented. After both intervals were presented, participants were asked to indicate which of the two intervals contained the flicker – this response was untimed. Intervals were indicated with an auditory cue. Participants could take a variable-length break in between blocks. Block-order was pseudorandom. **b)** Control results. Normalized cone-directed flicker detection thresholds on low- and high-LMS backgrounds, for the three participants in Experiment 1. Dark gray lines express thresholds as Just-Noticeable-Difference (JND) from the background on four single sessions. Solid blue lines indicate the median JNDs for each participant across four sessions. All JNDs for a participant were normalized to that participant's median JND on the low-melanopic background. Error bars indicate +/- 1 standard error of the median. If the cone-directed flicker detection is mediated by background LMS stimulation in a Weber's law fashion, JNDs on the 4.5x higher LMS background should be 4.5x higher (difference of 3.5 in the normalized JND representation) than on the low-LMS background (dashed horizontal line). Insets: The four red dots show the within-session differences between JNDs at low- and high-LMS adaptation level. The red horizontal line shows the median of the four within-session differences.
